# Spatial regulation of AMPK signaling revealed by a sensitive kinase activity reporter

**DOI:** 10.1101/2021.10.11.463987

**Authors:** Danielle L. Schmitt, Stephanie D. Curtis, Allen Leung, Jin-fan Zhang, Mingyuan Chen, Catherine Y. He, Sohum Mehta, Padmini Rangamani, Reuben J. Shaw, Jin Zhang

## Abstract

AMP-activated protein kinase (AMPK) is a master regulator of cellular energetics which coordinates metabolism by phosphorylating a plethora of substrates throughout the cell. But whether AMPK activity is regulated at different subcellular locations to provide precise spatial and temporal control over metabolism is unclear. Genetically encoded AMPK activity reporters (AMPKAR) have provided a window into spatial AMPK activity, but the limited dynamic range of current AMPKARs hinders detailed study. To monitor the dynamic activity of AMPK with high sensitivity, we developed a single-fluorophore AMPK activity reporter (ExRai AMPKAR) that exhibits an excitation ratiometric fluorescence change upon phosphorylation by AMPK, with over 3-fold greater response compared to previous AMPKARs. Using subcellularly localized ExRai AMPKAR, we found that the activity of AMPK at the lysosome and mitochondria are differentially regulated. While different activating conditions, irrespective of their effects on ATP, robustly yet gradually increase mitochondrial AMPK activity, lysosomal AMPK activity accumulates with much faster kinetics. Genetic deletion of the canonical upstream kinase liver kinase B1 (LKB1) resulted in slower AMPK activity at lysosomes but did not affect the response amplitude at either location, in sharp contrast to the necessity of LKB1 for maximal cytoplasmic AMPK activity. We further discovered AMPK activity in the nucleus, which resulted from LKB1-mediated cytoplasmic activation of AMPK followed by nuclear shuttling. Thus, a new, sensitive reporter for AMPK activity, ExRai AMPKAR, in complement with mathematical and biophysical methods, captured subcellular AMPK activity dynamics in living cells and unveiled complex regulation of AMPK signaling within subcellular compartments.

## Introduction

AMP-activated protein kinase (AMPK) is a ubiquitously expressed heterotrimeric protein in mammals, composed of one of two α kinase subunits, one of two β regulatory subunits, and one of three γ nucleotide-binding subunits^1^. Allosteric activation of AMPK is achieved through binding of adenine nucleotides to the γ subunit or small molecules to the allosteric drug and metabolite (ADAM) site at the interface between α and β subunits^2,3^. AMPK is regulated by several upstream kinases, predominantly liver kinase B1 (LKB1)^4,5^ and calcium/calmodulin protein kinase kinase 2 (CaMKK2)^6^, to control metabolic processes, including glycolysis, lipid and protein biosynthesis, mitochondrial biogenesis, and gene expression^7,8^.

How AMPK senses different stimuli and relays diverse signals to downstream components in various subcellular locations with high specificity has remained elusive. An emerging view is that compartmentalized AMPK signaling enables specificity towards downstream effectors^9,10^. Signaling complexes containing AMPK are found at different subcellular locations, including lysosomes, mitochondria, endoplasmic reticulum, and the nucleus^7^. Recent studies found that lysosomal pools of AMPK are preferentially activated by glucose starvation, whereas more severe metabolic stressors like glutamine starvation or pharmacological stimulation are required to activate cytosolic and mitochondrial pools of AMPK, due to specific assembly of lysosomal AMPK-activating signaling complexes^11^. However, our understanding of the regulation of AMPK signaling at different subcellular locations is still limited, and AMPK regulation in specific locations remains controversial^9,10^.

Genetically encoded reporters are powerful tools for interrogating the precise spatiotemporal regulation of signaling pathways^12,13^. Addition of localization tags for cellular organelles and compartments to kinase activity reporters enables subcellular spatial resolution of kinase activities in single cells, allowing for the elucidation of compartment-specific signaling mechanisms^14,15^. This approach enabled us to profile subcellular AMPK activity using a Förster Resonance Energy Transfer (FRET)-based AMPK activity reporter^16^. However, the dynamic range of the current FRET-based AMPK activity reporters^15–19^ limited our ability to fully distinguish the dynamics and regulation of spatiotemporal AMPK activity.

In this work, we developed a single-fluorophore excitation-ratiometric AMPK activity reporter (ExRai AMPKAR), which reports endogenous AMPK activity in living cells with high sensitivity. Using our new reporter alongside mathematical modeling, we found mitochondrial AMPK activity robustly responds to energy stress. We profiled spatiotemporal AMPK activity and found spatially defined roles for LKB1 in regulating cytoplasmic, lysosomal, and mitochondrial AMPK activity. Finally, we measured nuclear AMPK activity with ExRai AMPKAR and identified a regulatory mechanism of nuclear AMPK activity in living cells. This work provides a better understanding of spatiotemporal AMPK activity, which is critical for the diverse signaling profile of AMPK throughout the cell.

## Results

### Development of an excitation-ratiometric AMPK activity reporter

Building on our recent success in engineering single-fluorophore-based kinase activity reporters with high dynamic range^20,21^, we set out to develop a single-fluorophore AMPKAR. The modular design of biosensors enables coupling of different activity sensing units with readout-generating reporting units to create new biosensors^22,23^. We inserted a circularly permutated enhanced green fluorescent protein (cpEGFP), a widely used reporting unit^24^, between the two components of the AMPK activity sensing unit from a previous FRET-based AMPK biosensor^18^, which consists of an AMPK substrate domain^17^ and the phosphoamino acid-binding forkhead associated domain 1 (FHA1). Upon phosphorylation of the reporter and binding of the FHA1 domain to the phosphorylated AMPK substrate sequence, we expect to observe an increase in fluorescence emission with 488 nm excitation and a decrease with 400 nm excitation (**Fig 1a**). The ratio of these two fluorescence intensities (Ex 488 nm/400 nm) can be used as an excitation ratiometric readout (R). Given that linkers at the junctions of the sensing unit and reporting unit are particularly important in determining the dynamic range of the reporter^25^, we set out to vary the two amino acids immediately flanking either side of cpEGFP to identify a linker combination that would result in a large excitation ratio change upon AMPK stimulation. The top six linkers from our previous screen to generate a high-performance excitation-ratiometric protein kinase A activity reporter were selected^21^, and we tested the linker variants in Cos7 cells stimulated with an AMPK activator, 2-deoxyglucose (2-DG, **Supplemental Fig 1a**). From this linker screen, we found the linker combination FC/LL yielded the largest maximum ratio change (ΔR/R_0_), and this variant was selected for further characterization.

**Fig 1.**
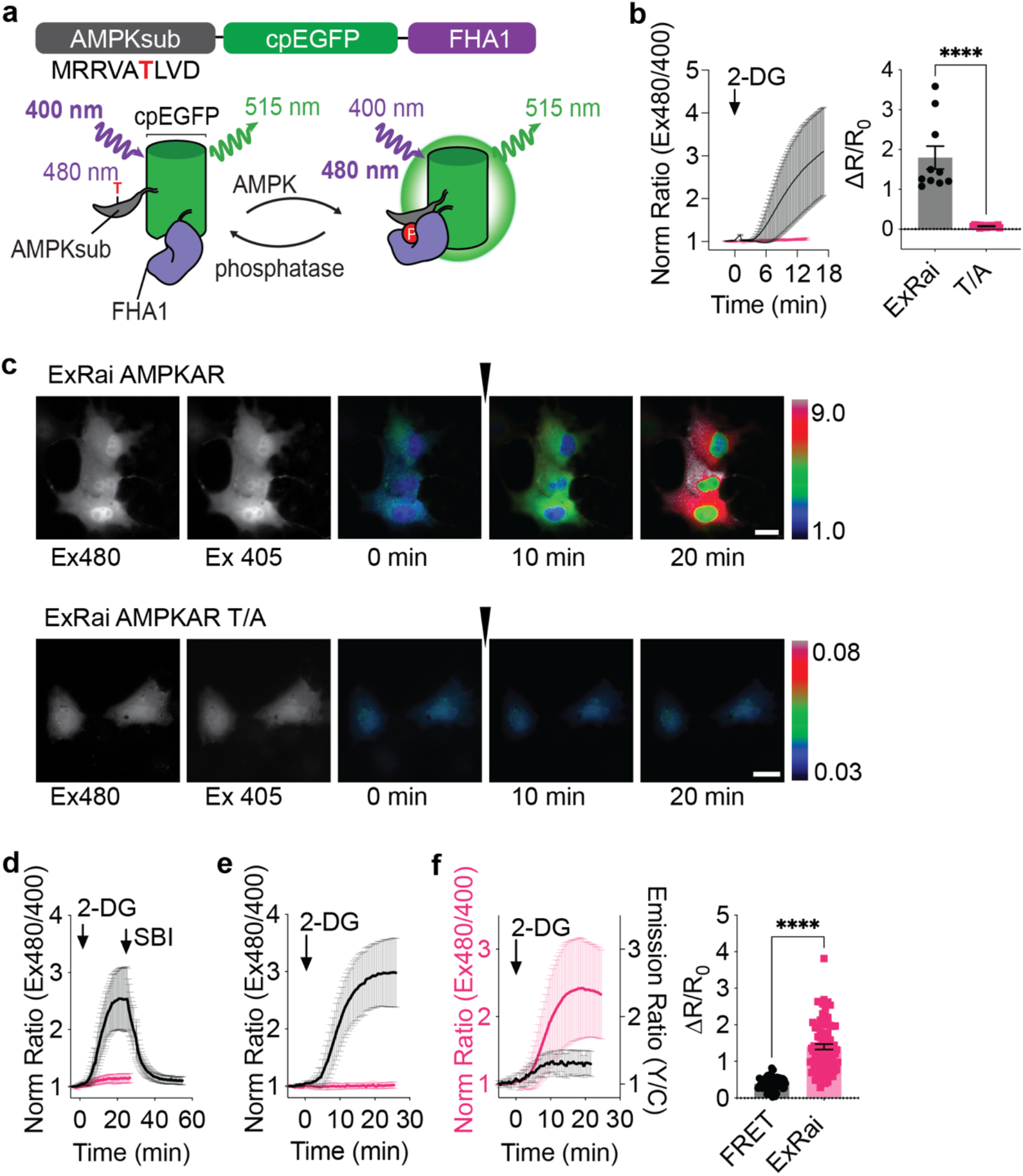
Development and characterization of ExRai AMPKAR. **a**, Design and domain structure of ExRai AMPKAR. Threonine phosphorylated by AMPK denoted in red. **b**, Average response of ExRai AMPKAR (black, n = 10 cells from 3 experiments), and ExRai AMPKAR T/A (pink, n = 17 cells from 3 experiments) to 2-DG (40 mM) stimulation in Cos7 cells along with maximum ratio change (****P<0.0001, unpaired t-test). **c**, Representative images of ExRai AMPKAR (top) and ExRai AMPKAR T/A (bottom) in Cos7 cells treated with 2-DG (40 mM) at the indicated time. **d,** Average response of ExRai AMPKAR to AMPK stimulation by 2-DG (40 mM) followed by SBI-0206965 (30 μM, black, n = 35 cells from 3 experiments) or after pretreatment with SBI-0206965 (pink, n = 43 cells from 4 experiments). **e**, Average response of ExRai AMPKAR in WT MEFs (black, n = 31 cells from 4 experiments) and AMPKα KO MEFs (pink, n = 23 cells from 4 experiments) treated with 2-DG. **f**, Response of ExRai AMPKAR (pink, n = 90 cells from 5 experiments) and ABKAR (black trace, n = 25 cells from 2 experiments) in HEK293T cells treated with 2-DG (40 mM). For all figures, time courses show the mean ± SD, dot plots show the mean ± SEM. Scale bars, 20 μm.

Stimulation of AMPK using 2-DG in Cos7 cells expressing this reporter led to a large change in the excitation ratio (ΔR/R_0_ = 1.80 ± 0.29; **Fig 1b-c, Supplemental Fig 1b**), whereas mutating the phosphorylation site threonine to alanine (T/A) to generate a phospho-null mutant resulted in a biosensor with minimal response (ΔR/R_0_ = 0.075 ± 0.0064, P < 0.0001). We tested the reversibility of the response using the AMPK inhibitor SBI-0206965 (SBI)^26,27^. In HEK293T cells expressing the reporter construct, 2-DG-stimulated AMPK activity was rapidly suppressed by SBI-0206965 to near basal levels (R = 1.11 ± 0.011; **Fig 1d**). Similarly, pretreatment with SBI-0206965 blocked the 2-DG-induced reporter response (ΔR/R_0_ = 0.17 ± 0.011). To confirm the reporter responses are specific for AMPK activity, we expressed our biosensor in either wild-type or AMPKα1/2 knockout mouse embryonic fibroblasts^28^ (WT or AMPKα KO MEFs, **Supplemental Fig 1c**). MEFs were then treated with 2-DG, which induced large excitation-ratio changes in WT MEFs, whereas minimal responses were observed in AMPKα KO MEFs (**Fig 1e**).

Finally, we compared our new construct to our previous FRET-based reporter^18^. In HEK293T cells stimulated with 2-DG, the excitation-ratio change from our new construct (ΔR/R_0_ = 1.40 ± 0.072) was larger than the yellow/cyan emission ratio change from the FRET-based biosensor (ΔR/R_0_ = 0.39 ± 0.024, P < 0.0001; **Fig 1f**), which translated to a higher signal-to-noise ratio (SNR; FRET biosensor: 59.11 ± 6.46 vs new construct: 228.1 ± 12.81, P < 0.0001, **Supplementary Fig 1d**). We also estimated the Z-factor, a measurement of assay fitness^29^. From the average responses of these two reporters, the FRET-based biosensor had a Z-factor of 0.44, while our new construct had a Z-factor of 0.90, indicating better assay fitness. Thus, we have developed a new AMPK activity reporter which we termed excitation-ratiometric AMPKAR (ExRai AMPKAR), for specific monitoring of dynamic AMPK activity within living cells with greatly enhanced sensitivity.

### Mitochondrial AMPK activity robustly responds to energy stresses irrespective of their effects on ATP

AMPK regulates mitochondrial fission and mitophagy, and we have previously reported that AMPK is active at mitochondria^16,28,30^. However, how mitochondrial AMPK activity responds to different metabolic stressors is not clear. To better visualize dynamics of mitochondrial AMPK activity, we fused ExRai AMPKAR to a mitochondria-targeting sequence from Dual specific A-Kinase Anchoring Protein 1 (DAKAP1) and expressed the targeted reporter in either WT or AMPKα KO MEFs (**Fig 2a**). WT MEFs expressing mitochondrial-localized ExRai AMPKAR and treated with 2-DG showed strong mitochondrial AMPK activity, while AMPKα KO MEFs exhibited minimal AMPK activity (**Fig 2b**).

**Fig 2.**
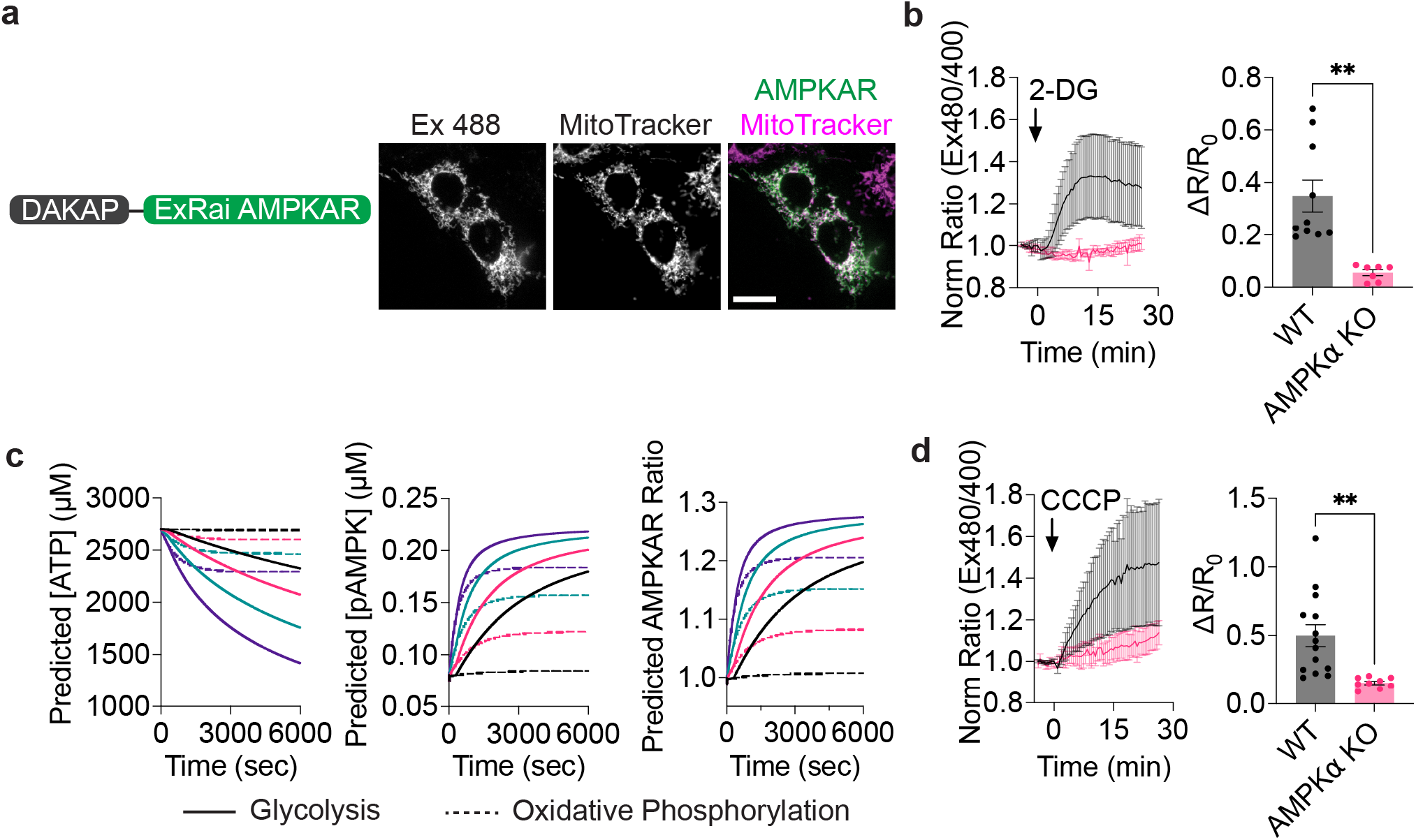
Mitochondrial AMPK activity robustly senses metabolic stress. **a**, Domain layout and representative image of mito-ExRai AMPKAR in MEFs stained with the mitochondrial marker MitoTracker Red. **b**, Average response of mito-ExRai AMPKAR in WT (black, n = 10 cells from 5 experiments and AMPKα KO MEFs (pink, n = 7 cells from 3 experiments) treated with 2-DG (40 mM) along with maximum ratio change (**P = 0.0012, unpaired t-test). **c**, Predicted ATP concentration, pAMPK concentration, and AMPKAR response across a range of ATP hydrolysis rates (0.5 μM/s black, 2 μM/s pink, 4 μM/s teal, 8 μM/s purple; solid lines indicate inhibition of glycolysis, dashed lines indicate inhibition of oxidative phosphorylation). **d**, Average response of mito-ExRai AMPKAR in WT (black, n = 14 cells from 5 experiments) and AMPKα KO MEFs (pink, n = 9 cells from 3 experiments) upon CCCP (5 μM) stimulation, along with maximum ratio change (**P = 0.0027, unpaired t-test). For all figures, time courses show the mean ± SD, dot plots show the mean ± SEM. Scale bars, 20 μm.

To better understand how mitochondrial AMPK activity is influenced by energy stress, we sought to use our single-cell measurements of metabolic stress-induced AMPK activity to develop a mathematical model of mitochondrial AMPK activity to predict how AMPK responds to different metabolic stresses. Our mathematical model was influenced by other approaches which incorporate biosensor measurements and adenine nucleotide equilibrium^31,32^. This enabled mitochondrial-localized ExRai AMPKAR measurements to be modeled as Michaelis-Menten kinetics and metabolic fluxes as mass-action kinetics (**Supplemental Tables 1-3**). We then scaled the rates of ATP hydrolysis for either glycolysis or oxidative phosphorylation to assess the impact of each pathway on ATP concentration, phosphorylated AMPK (pAMPK), and ExRai AMPKAR output. By doing this, we can model how discreet changes in ATP production through inhibition of glycolysis or oxidative phosphorylation impact AMPK activity at the mitochondria. Across a range of ATP hydrolysis rates (0.5-8 μM/s) simulating inhibition of either glycolytic or oxidative ATP production, mitochondrial AMPK activity robustly responded to metabolic stress (**Fig 2c**). While inhibition of glycolysis was more effective than inhibition of oxidative phosphorylation at depleting ATP, likely due to the cascading effect of glycolytic inhibition to deplete ATP production through oxidative phosphorylation, even small perturbations in either glycolysis or oxidative phosphorylation were predicted to strongly induce AMPK activity.

To test the model prediction that AMPK robustly responds to energy stress, regardless of source, we activated AMPK in MEFs using carbonyl cyanide m-chlorophenyl hydrazone (CCCP), an uncoupler of mitochondrial oxidative phosphorylation, which has been shown to activate AMPK in the mitochondrial fraction^30^. CCCP induced mitochondrial AMPK activity to similar levels as 2-DG treatment in living cells, as measured by ExRai AMPKAR (2-DG: ΔR/R_0_ = 0.35 ± 0.061; CCCP: ΔR/R_0_ = 0.50 ± 0.080, P = 0.24; **Fig 2d**), consistent with the model prediction. By combining mathematical modeling with live-cell imaging, we find that mitochondrial AMPK activity robustly responds to energy stresses.

### Rapid AMPK activity at the lysosome is dependent on LKB1

The mitochondria and lysosome represent key signaling locations for AMPK^11,28,33–37^. Notably, results from western blotting assays have indicated that lysosomal pools of AMPK are preferentially activated under glucose deprivation, due to a lysosomal-localized AMPK regulatory complex^38^, while mitochondrial AMPK is activated under more severe nutrient stress^11^. This suggests the lysosome could serve as a signaling hub for AMPK. To specifically investigate lysosomal AMPK activity, we fused ExRai AMPKAR to lysosome associated membrane protein 1 (LAMP1, **Fig 3a**), and WT and AMPKα KO MEFs expressing lysosomal-targeted ExRai AMPKAR were treated with 2-DG (**Fig 3b**). We then compared the time-to-half-maximum (t_1/2_) of cytoplasmic, mitochondrial, and lysosomal ExRai AMPKAR responses to 2-DG to identify kinetic differences amongst these locations. Lysosomal activity increased more rapidly than either cytoplasmic or mitochondrial activity (t_1/2_ = 3.09 ± 0.30 min, P < 0.0001; **Fig 3c**). Therefore, we sought to identify the mechanism underlying rapid accumulation of AMPK activity at the lysosome.

**Fig 3.**
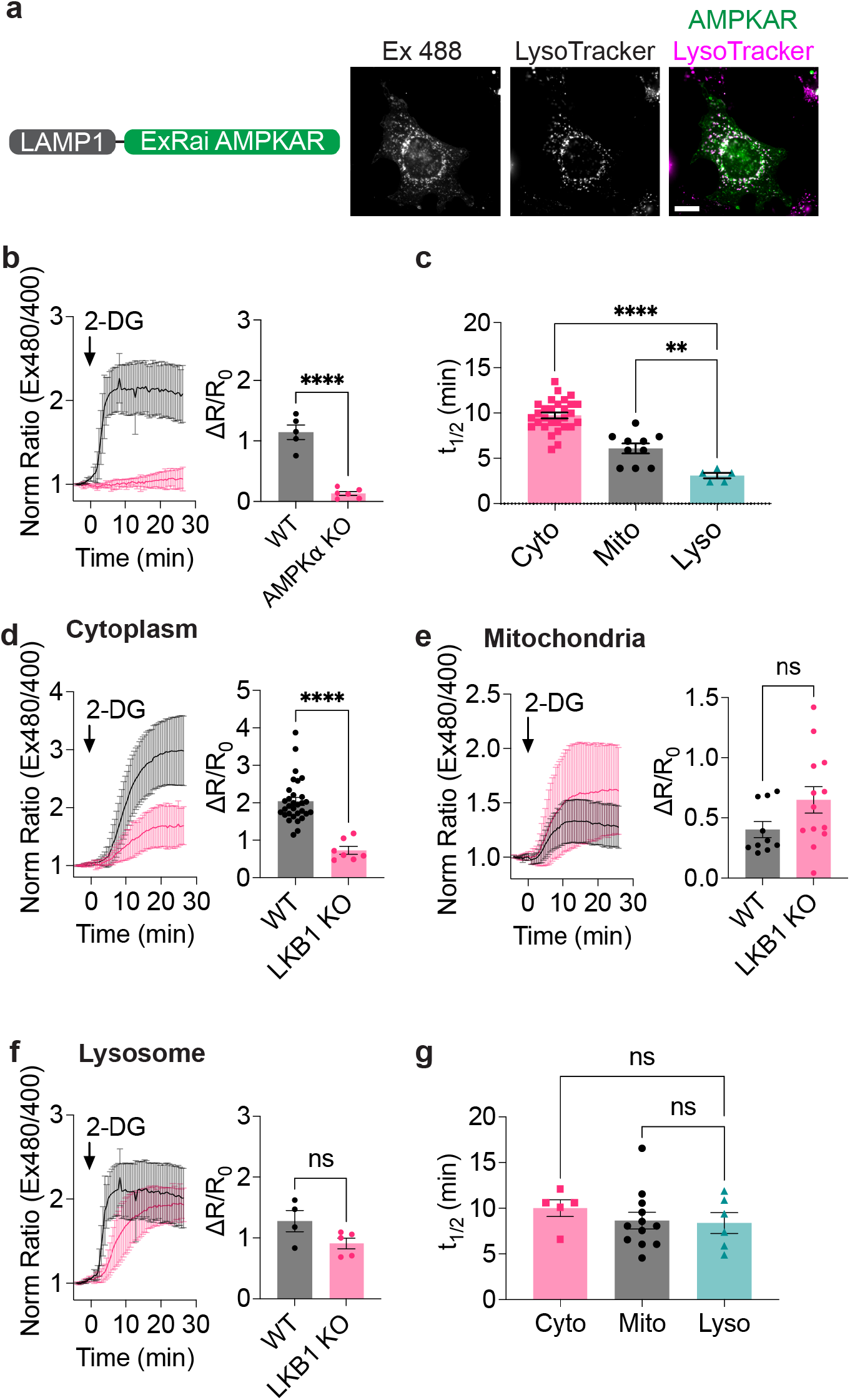
Lysosomal AMPK activity is rapidly induced in an LKB1-dependent manner. **a**, Domain layout and representative image of lyso-ExRai AMPKAR in MEFs stained with the lysosomal marker LysoTracker Red. **b**, Average response of lyso-ExRai AMPKAR in WT (black, n = 5 cells from 2 experiments) and AMPKα KO MEFs (pink, n = 6 cells from 2 experiments) to 2-DG (40 mM) stimulation, along with maximum ratio change (P<0.0001, unpaired t-test). **c**, Time to half-maximal response (t_1/2_) of cytoplasmic ExRai AMPKAR (pink), mito-ExRai AMPKAR (black), and lyso-ExRai AMPKAR (teal) following treatment with 2-DG (**P = 0.0033; ****P<0.0001, one-way ANOVA with Dunnett’s multiple comparisons test). **d**, Average response to 2-DG (40 mM) stimulation of cytoplasmic ExRai AMPKAR in WT (black, reproduced from Fig 1e) and LKB1 KO MEFs (pink, n = 7 cells from 4 experiments), along with maximum ratio changes (****P<0.0001) **e**, Average response to 2-DG (40 mM) stimulation of mito-ExRai AMPKAR in WT (black, reproduced from Fig 2b) and LKB1 KO MEFs (pink, n = 13 cells from 5 experiments), along with maximum ratio changes (ns P > 0.085, unpaired t-test). **f**, Average response to 2-DG (40 mM) stimulation of lyso-ExRai AMPKAR in WT (black, reproduced from Fig 3b) and LKB1 KO MEFs (pink, n = 6 cells from 4 experiments), along with maximum ratio changes (ns P > 0.085, unpaired t-test). **g**, t_1/2_ of cytoplasmic ExRai AMPKAR (pink), mito-ExRai AMPKAR (black), and lyso-ExRai AMPKAR (teal) following treatment with 2-DG in LKB1 KO MEFs (ns P > 0.55, one-way ANOVA with Dunnett’s multiple comparisons test). For all figures, time courses show the mean ± SD, dot plots show the mean ± SEM. Scale bars, 20 μm.

Lysosomal regulation of AMPK activity is thought to occur via a multi-protein complex which coordinates the lysosomal localization of AMPK and its upstream kinase LKB1^34,35^. We hypothesized that the compartmentation of LKB1 and AMPK together at the lysosome is responsible for the fast kinetics of lysosomal AMPK activity compared to other locations. We therefore compared AMPK activity at the cytoplasm, mitochondria, and lysosome in WT and LKB1 KO MEFs^39^ (**Supplemental Fig 2a**). 2-DG-induced cytoplasmic AMPK activity was significantly suppressed in LKB1 KO MEFs compared to WT MEFs, consistent with the established prominent role of LKB1 in mediating 2-DG-induced AMPK activity^40^ (2-DG ΔR/R_0_ WT MEF 2.04 ± 0.10; LKB1 KO MEF 0.73 ± 0.11, P<0.0001; **Fig 3d**). At the mitochondria, the absence of LKB1 had minimal effect on mitochondrial AMPK activity in response to 2-DG (**Fig 3e and Supplemental Fig 2b**). At the lysosome, we found that while 2-DG could still induce AMPK activity in LKB1 KO MEFs, the kinetics were substantially slower than in WT MEFs (t_1/2_ 8.39 ± 1.14 min, P = 0.0026; **Fig 3f**). The kinetics of AMPK activity at the cytoplasm, mitochondria and lysosome were similar in LKB1 KO MEFs (**Fig 3g**), indicating that LKB1 KO eliminated the kinetic advantage of lysosomal AMPK activity. These results suggest that the presence of LKB1 drives the fast kinetics of AMPK activity at the lysosome, most likely through close localization of LKB1 and AMPK at this organelle. Taken together, our findings reveal spatially distinct regulatory control of LKB1 over AMPK activity.

### AMPK activity induced by allosteric activators exhibits a distinct spatial profile

Allosteric activation of AMPK, independent of energy stress or calcium signaling, is an attractive therapeutic route for treating diabetes and other metabolic disorders^41^. Towards this goal, several synthetic ligands which bind the ADAM site have been designed^42–44^, including MK-8722^45,46^. Recent work has also identified endogenous ligands for the ADAM site^47^. However, how ADAM site activators influence spatially compartmentalized AMPK signaling is not clear. Mechanistically, the importance of upstream kinases for ADAM site activation remains controversial^48,49^, as ADAM site activators protect AMPK from dephosphorylation by phosphatases^3,48,50^. As kinase and phosphatase activities can vary based on location^12,13^, we hypothesized there could be spatially distinct differences in allosteric activator induced AMPK activity. To test this hypothesis, we set out to use ExRai AMPKAR to profile spatiotemporal AMPK activity in response to allosteric activation via the ADAM site and determine the importance of the upstream kinase LKB1 for spatially defined, allosterically activated AMPK.

We first measured MK-8722-induced cytoplasmic AMPK activity in WT and AMPKα KO MEFs, finding MK-8722 strongly induced AMPK activity in the cytoplasm (**Fig 4a**). We assessed the necessity of LKB1 for MK-8722-mediated AMPK activity. In LKB1 KO MEFs, MK-8722 induced cytoplasmic AMPK activity was significantly reduced compared with WT MEFs (WT: ΔR/R_0_ = 1.57 ± 0.16 vs LKB1: ΔR/R_0_ = 0.66 ± 0.12, P = 0.0041; **Fig 4a**). These data suggest that LKB1 is required for the maximal cytoplasmic AMPK activity induced by the allosteric activator MK-8722.

**Fig 4.**
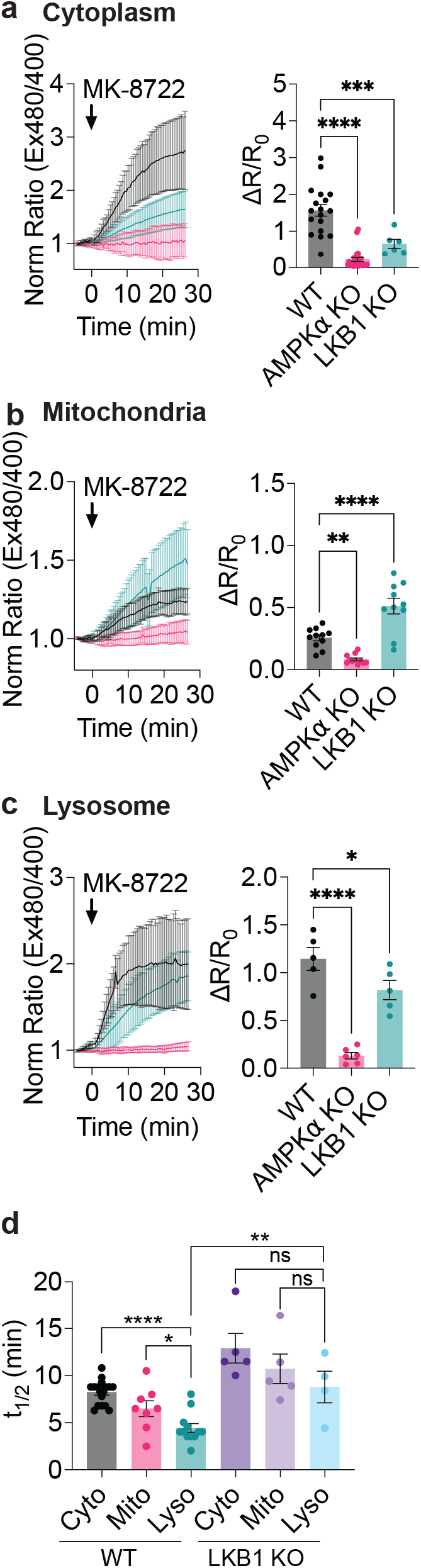
AMPK activity induced by MK-8722 exhibits distinct spatiotemporal dynamics. **a**, Average response of ExRai AMPKAR in WT MEFs (black, n = 19 cells from 3 experiments), AMPKα KO (pink, n = 25 cells from 3 experiments), and LKB1 KO MEFs (teal, n = 6 cells from 4 experiments) treated with MK-8722 (500 nM) along with maximum ratio change (***P = 0.002, ****P<0.0001, one-way ANOVA with Dunnett’s multiple comparisons test). **b**, Average response of mito-ExRai AMPKAR in WT (black, n = 11 cells from 4 experiments), AMPKα KO (pink, n = 11 cells from 3 experiments), and LKB1 KO MEFs (teal, n = 10 cells from 5 experiments) treated with MK-8722 (500 nM) along with maximum ratio change (**P = 0.0048, ****P<0.0001, one-way ANOVA with Dunnett’s multiple comparisons test). **c**, Average response of lyso-ExRai AMPKAR in WT (black, n = 12 cells from 3 experiments), AMPKα KO (pink, n = 8 cells from 3 experiments) and LKB1 KO MEFs (teal, n = 5 cells from 4 experiments) to MK-8722 (500 nM) stimulation, along with maximum ratio change (*P = 0.041, ***P = 0.0002, one-way ANOVA with Dunnett’s multiple comparisons test). **d**, t_1/2_ of cytoplasmic ExRai AMPKAR (black), mito-ExRai AMPKAR (pink), and lyso-ExRai AMPKAR (teal) expressed in WT MEFs and cytoplasmic ExRai AMPKAR (dark purple), mito-ExRai AMPKAR (light purple), and lyso-ExRai AMPKAR (light blue) expressed in LKB1 KO MEFs following treatment with MK-8722. (*P = 0.0203; ****P<0.0001, ns > 0.05, one-way ANOVA with Dunnett’s multiple comparisons test, **P = 0.0031, unpaired t-test). For all figures, time courses show the mean ± SD, dot plots show the mean ± SEM.

Next, we examined mitochondrial and lysosomal AMPK activity induced by MK-8722. WT and AMPKα KO MEFs expressing mitochondrial- or lysosomal-targeted ExRai AMPKAR were treated with MK-8722 (**Fig 4b-c**). MK-8722 robustly induced AMPK activity at these locations. We found lysosomal AMPK activity was more rapid than either mitochondrial or cytoplasmic AMPK activity (4.44 ± 0.47 min, P < 0.0001; **Fig 4d**), suggesting that there are spatially distinct differences in AMPK activity induced by allosteric activators.

LKB1 KO also showed differential effect on MK-2733 induced subcellular AMPK activities. In LKB1 KO cells expressing lysosomal-targeted ExRai AMPKAR, we found MK-8722 still induced strong lysosomal AMPK activity, but with slower kinetics compared to WT MEFs, like that observed with 2-DG treatment (t_1/2_ 8.79 ± 1.68 min, P = 0.0031; **Fig 4c-d**). On the other hand, the absence of LKB1 had minimal effect on mitochondrial AMPK activity in response to MK-8722 (**Fig 4b and Supplemental Fig 2c**), suggesting allosterically induced mitochondrial AMPK activity does not require an upstream kinase or another basally active upstream kinase is present at the mitochondria^19^. Together, we found the effect of ADAM site activation is not only dependent on AMPK and ADAM site ligands, but also relies on regulatory mechanisms, like upstream kinases, that are embedded in the spatiotemporal signaling network.

### Nuclear AMPK activity measured using ExRai AMPKAR

AMPK has been implicated in the regulation of various nuclear targets^8,9^, and AMPK subunits have been reported to directly localize to the nucleus^51^. Nevertheless, the existence and regulation of nuclear AMPK activity remain controversial^9,10^. Biosensor-based studies have thus far been unable to conclusively shed light on the presence of nuclear activity^16,17,52^, with some studies suggesting that AMPK is active in the nucleus in response to energy stress and others failing to detect any nuclear AMPK activity^11,53–56^. We therefore hypothesized that the enhanced sensitivity of ExRai AMPKAR would allow us to provide a more definitive view of nuclear AMPK activity.

ExRai AMPKAR was fused to a nuclear localization sequence (NLS), and we confirmed ExRai AMPKAR nuclear localization (**Fig 5a**). In WT MEFs, we observed significant AMPK activity in the nucleus following treatment with 2-DG, with minimal activity detected in AMPKα KO MEFs (WT: ΔR/R_0_ = 0.55 ± 0.05 vs AMPKα KO: ΔR/R_0_ = 0.11 ± 0.013, P < 0.0001; **Fig 5b**). Similarly, MK-8722 induced nuclear AMPK activity in WT MEFs, whereas AMPKα KO MEFs showed minimal responses from nuclear localized ExRai AMPKAR (WT: ΔR/R_0_ = 0.43 ± 0.030 vs AMPKα KO: ΔR/R_0_ = 0.15 ± 0.026, P < 0.001; **Fig 5c**). In parallel, we directly probed for phosphorylated AMPKα in the nuclei of WT MEFs using nuclear fractionation. Cells were treated with DMSO, 2-DG, or MK-8722, and nuclear fractions collected and probed for phosphorylated AMPKα via western blotting. We found that along with cytoplasmic pools of AMPKα becoming phosphorylated following treatment with 2-DG or MK-8722, nuclear pools of AMPK were phosphorylated as well (**Fig 5d**). Thus, our results provide clear evidence of nuclear AMPK activity induced by both metabolic stress and allosteric activation.

**Fig 5.**
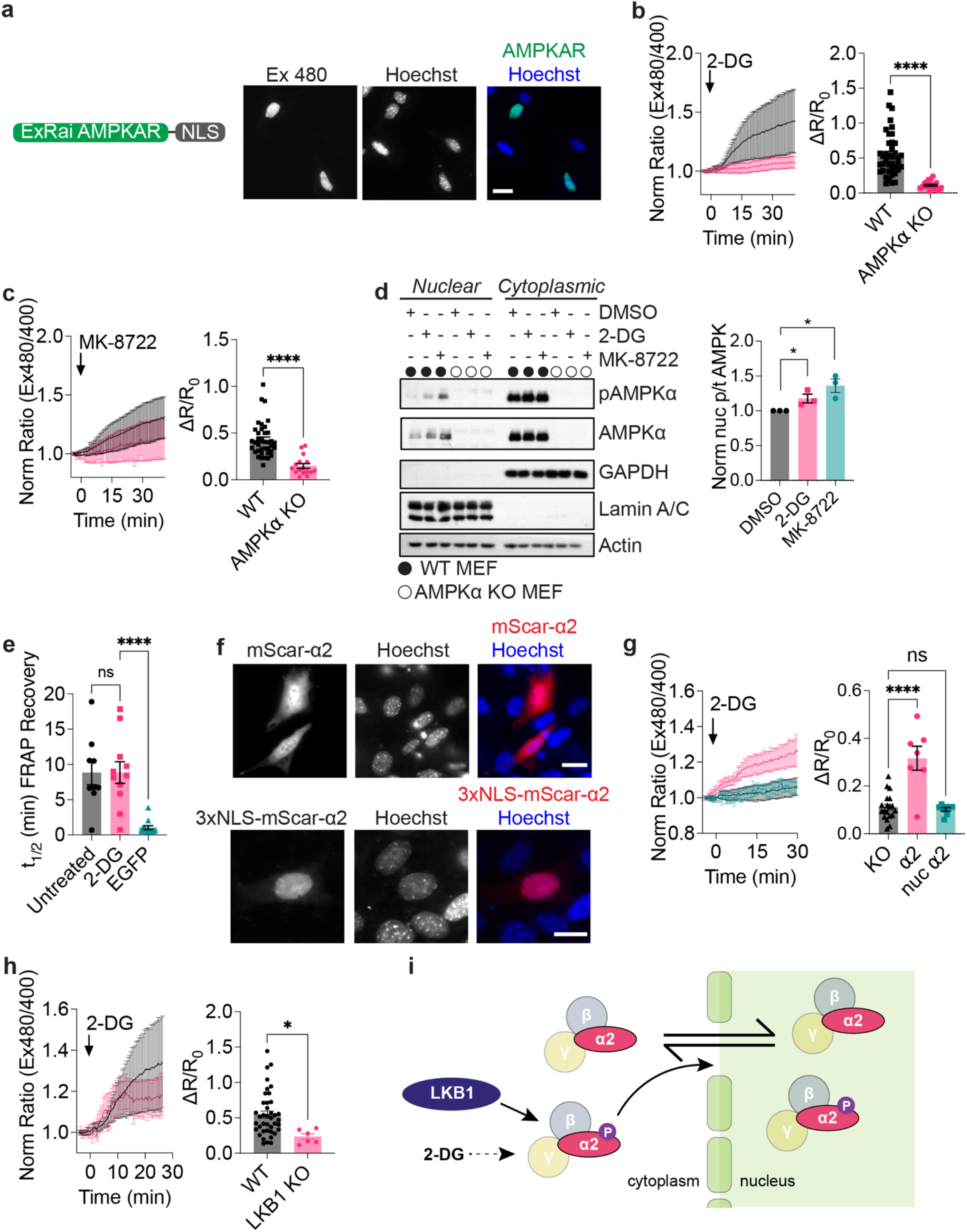
Nuclear AMPK activity measured using ExRai AMPKAR. **a**, Domain layout and representative image of ExRai AMPKAR-NLS expressed in MEFs stained with Hoechst nuclear marker. **b**, Average response of ExRai AMPKAR-NLS in either WT (black, n = 46 cells from 5 experiments) or AMPKα KO MEFs (pink, n = 19 cells from 4 experiments) treated with 2-DG (40 mM), along with maximum ratio change (****P < 0.0001, unpaired t-test). **c**, Average response of ExRai AMPKAR-NLS in either WT (black, n = 38 cells from 9 experiments) or AMPKα KO MEFs (pink, n = 16 cells from 2 experiments) treated with MK-8722 (500 nM), along with maximum ratio change (****P < 0.0001, unpaired t-test). **d**, Western blot of nuclear-fractionated MEFs treated with DMSO, 2-DG (40 mM), or MK-8722 (500 nM) for 60 min. Quantification from three independent trials (*P < 0.05, unpaired t-test). **e**, Half-time of FRAP recovery (min) for nuclear EGFP-AMPKα2 in AMPKα KO MEFs either without (n = 9 cells from 3 experiments) or with 2-DG stimulation (40 mM, n = 11 cells from 3 experiments) immediately before FRAP experiment began, or for EGFP alone (n = 13 cells from 2 experiments). (****P < 0.0001, one-way ANOVA with Dunnett’s multiple comparisons test). **f**, Representative images of mScarlet-AMPKα2 or 3xNLS-mScarlet-AMPKα2 in MEFs stained with Hoechst nuclear marker. **g**, Average 2-DG (40 mM)-stimulated response of AMPKα KO MEFS expressing ExRai AMPKAR-NLS alone (black, reproduced from Fig 4b) or co-expressing mScarlet-AMPKα2 (pink, n = 7 cells from 2 experiments) or 3x NLS-mScarlet-AMPKα2 (teal, n = 8 cells from 4 experiments), along with maximum ratio change (****P < 0.0001, one-way ANOVA with Dunnett’s multiple comparisons test). **h**, Average response of ExRai AMPKAR-NLS in either WT (black, reproduced from Fig 4b) or LKB1 KO MEFs (pink, n = 6 cells from 4 experiments) treated with 2-DG (40 mM), along with maximum ratio change (*P = 0.021, unpaired t-test). **i**, Mechanism of 2-DG-induced nuclear AMPK activity where nuclear AMPK activity in response to 2-DG is initiated in the cytoplasm in an LKB1-dependent manner, after which AMPK then translocates into the nucleus to phosphorylate nuclear targets. For all figures, time courses show the mean ± SD, dot plots show the mean ± SEM. Scale bars, 20 μm.

### Nuclear AMPK activity requires AMPKα2 nucleo-cytoplasmic shuttling

We next investigated the mechanism behind 2-DG-induced nuclear AMPK activity. We considered two possible scenarios for nuclear AMPK activity. The first scenario considers AMPK to reside within the nucleus and become activated *in situ*. In the second scenario, we considered AMPK to shuttle between the cytoplasm and nucleus, with AMPK activity being initiated in the cytoplasm, followed by translocation of active AMPK into the nucleus. To differentiate between these two possibilities, we first sought to determine if AMPK translocates into the nucleus under basal and stimulated conditions. For this purpose, we focused on AMPKα2, which has been detected in both the cytoplasm and nucleus^51^. To measure AMPK translocation, we performed fluorescence recovery after photobleaching (FRAP) experiments and monitored the nuclear intensity of EGFP-tagged AMPKα2 (EGFP-AMPKα2) in AMPKα KO MEFs after photobleaching of the nuclei. Under basal conditions, nuclear EGFP-AMPKα2 intensity was fully recovered within 30 min (t_1/2_ = 8.84 ± 1.72 min; **Fig 5e, Supplemental Fig 3a**). EGFP-AMPKα2 nuclear intensity showed similar recovery kinetics following treatment with 2-DG (t_1/2_ = 8.86 ± 1.53 min; P = 0.999), suggesting that acute AMPK activation does not alter the kinetics of nuclear shuttling. Control cells expressing EGFP alone showed full recovery of nuclear intensity within only a few minutes (t_1/2_ = 1.06 ± 0.26 min, P<0.0001), indicating that the slower recovery time for nuclear EGFP-AMPKα2 fluorescence intensity is due to AMPKα2 itself and not the EGFP tag.

We then compared the t_1/2_ of EGFP-AMPKα2 import into the nucleus to that of 2-DG-induced nuclear AMPK activity. 2-DG induced nuclear AMPK activity within 15.53 ± 1.14 min, which is slower than the half-time for EGFP-AMPKα2 translocation into the nucleus. This supports our second scenario, whereby activated AMPK can translocate from the cytoplasm to the nucleus to induce nuclear signaling. To test if nuclear AMPK activity is dependent on AMPK shuttling into the nucleus, we sought to localize AMPKα to the nucleus and measure nuclear AMPK activity in AMPKα KO MEFs. To sequester AMPK in the nucleus, we considered known nuclear localization and exclusion sequences present in AMPKα2^57,58^. However, mutating either the nuclear exclusion or localization sequence on AMPKα2 yielded no significant effect on the nuclear localization of EGFP-tagged AMPKα2 in HEK293T cells (**Supplemental Fig 3b**). Indeed, the effects of these sequences on nuclear AMPK activity are thought to be cell-type specific. For example, the putative nuclear localization signal was identified in mouse myoblasts and found to be leptin-dependent over the course of several hours, and not functional in HEK293T cells^57,58^. Therefore, we set out to further understand how nuclear AMPK activity is regulated by using a subcellular targeting approach.

We generated a nuclear-targeted AMPKα2, 3xNLS-mScarlet-AMPKα2, with untargeted mScarlet-AMPKα2 as a control. While mScarlet-AMPKα2 can translocate from the cytoplasm to the nucleus, the addition of three tandem nuclear localization sequences effectively sequesters mScarlet-AMPKα2 in the nucleus (**Fig 5f**). We next examined the ability of these AMPKα2 constructs to rescue nuclear AMPK activity. Untargeted mScarlet-AMPKα2 restored nuclear AMPK activity following treatment with 2-DG in AMPKα KO MEFs (ΔR/R_0_ = 0.39 ± 0.063, P < 0.0001; **Fig 5g**), indicating that AMPKα2 is adequate for nuclear AMPK activity. However, expression of 3xNLS-mScarlet-AMPKα2 was insufficient to restore nuclear AMPK activity in AMPKα KO MEFs following treatment with 2-DG (ΔR/R_0_ = 0.10 ± 0.0075, P = 0.96). These results indicate that AMPK sequestered in the nucleus cannot respond to 2-DG stimulation and further support a mechanistic model of translocation-based nuclear AMPK activity. This suggests that nuclear AMPK activity will show a similar dependence on upstream kinases as cytoplasmic AMPK activity. Indeed, in LKB1 KO MEFs expressing nuclear ExRai AMPKAR, we found that nuclear AMPK activity was suppressed following 2-DG treatment (ΔR/R_0_ = 0.24 ± 0.04, P = 0.021; **Fig 5h**). We validated these results in HeLa cells with and without LKB1 expression (**Supplemental Fig 3c**). From our work, we propose a mechanistic model in which nuclear AMPK activity in response to 2-DG is initiated in the cytoplasm in an LKB1-dependent manner, after which AMPK then translocates into the nucleus to phosphorylate nuclear targets (**Fig 5i**).

## Discussion

Genetically encoded fluorescent protein-based kinase activity reporters have become essential tools to investigate compartmentalized signaling networks. In the present study, we designed and used a new single-fluorophore AMPK activity reporter, ExRai AMPKAR, the most sensitive AMPK activity reporter thus far developed. While FRET-based AMPKARs have been successfully used to interrogate AMPK activity^16–19^, the limited dynamic range of FRET-based reporters significantly limits their application. ExRai AMPKAR has over 3-fold higher dynamic range, enabling detection of subtle changes in AMPK activity. This allowed us to clearly detect nuclear AMPK activity, which has posed a challenge for FRET-based AMPKARs^16,17^. Therefore, ExRai AMPKAR now enables the robust visualization of subtle, subcellular AMPK signaling events.

Using ExRai AMPKAR, we interrogated the spatiotemporal dynamics of AMPK activity in response to two stimuli: energy stress and allosteric activation by ADAM site ligands. We found that lysosomal AMPK activity was more rapidly induced compared to mitochondrial or cytoplasmic AMPK activity. As the lysosome has been suggested to function as a signaling hub^38,41^, the rapid accumulation of AMPK activity at the lysosome could provide an advantage to substrates in proximity. Interestingly, we show that irrespective of the mode of activation, the kinetic trend for spatial AMPK activity is the same.

Building on these findings, we investigated the necessity of LKB1 for subcellular AMPK activity. At the lysosome, loss of LKB1 was associated with slower kinetics of AMPK activity, consistent with the demonstration of an AMPK regulatory complex at the lysosome consisting of AXIN, LKB1, and lysosomal membrane proteins such as vATPase and Ragulator^35^. While cytoplasmic AMPK activity was significantly diminished in the absence of LKB1, we found that LKB1 is not required for maximal lysosomal or mitochondrial AMPK activity. Consistent with our findings, AMPK activity has been detected in LKB1-deficient cells stimulated with 2-DG, which was abolished upon deletion or inhibition of CaMKK2^19^. 2-DG has been reported to increase intracellular calcium^19^, which along with AMP increases, could maximally activate AMPK through CaMKK2 in the absence of LKB1. As only cytoplasmic AMPK activity was significantly diminished in the absence of LKB1, it is possible the substituting upstream kinases, such as CaMKK2, exhibit enhanced activity around membranes. The necessity of upstream kinases for the effect of ADAM site activators is debated^48,49^, and our studies show location-specific dependence of MK-8722 on LKB1 to stimulate AMPK activity. While MK-8722 did not maximally induce AMPK activity in the cytoplasm in the absence of LKB1, impacts on MK-8722-induced AMPK activity at the mitochondria and lysosome were minimal. This could be due to the presence of other basally active upstream kinases or subcellular differences in phosphatase activities^59–61^, requiring further investigation.

With the increased sensitivity of ExRai AMPKAR, we report nuclear AMPK activity in response to treatment with 2-DG and MK-8722 and propose a mechanism for 2-DG-induced nuclear AMPK activity (**Fig 5i**). Basally, AMPKα2 can translocate between the cytoplasm and nucleus. As macromolecules larger than 40 kDa typically require the assistance of nuclear transport factors to cross the nuclear membrane^62^, future studies are needed to identify the molecular determinants responsible for shuttling AMPK, which comprises three subunits with a combined molecular weight > 100 kDa^63^. Our FRAP experiments suggested that the kinetics of AMPK nuclear shuttling are not affected by 2-DG. However, subcellular fractionation revealed slight increases in nuclear AMPK levels upon 2-DG stimulation (**Figure 5d**). Substantial translocation and accumulation of AMPKα subunits to the nucleus were reported after long-term treatment with leptin or adiponectin^32^, suggesting time- and stimulation-dependent accumulation of nuclear AMPK. Finally, we found that nuclear AMPK activity is reliant on LKB1. Others have reported that LKB1 is primarily active in the cytoplasm^64–66^, consistent with our model of LKB1 phosphorylating cytoplasmic AMPK, followed by nuclear translocation of phosphorylated AMPK to lead to subsequent nuclear activity. As our model suggests constant shuttling of AMPK between the cytoplasm and nucleus, it is possible that active AMPK is shuttled back to the cytoplasm, where it could be dephosphorylated. Taken together, our results present a new model of nuclear AMPK activity.

Our nuclear studies were limited to investigation of AMPKα2-containing complexes. We focused on AMPKα2 due to the known nuclear localization of this subunit^51^. However, emerging evidence suggests that AMPKα1 is vital for calcium-induced nuclear AMPK activity. Etoposide treatment specifically induced CaMKK2-dependent AMPK activity in the nucleus, but only with AMPKα1-containing complexes, even in the presence of AMPKα2 complexes^67^. CaMKK2-dependent AMPK activity has also been reported in response to a variety of physiological and pharmacological agents which increase intracellular calcium^68–73^, also dependent on AMPKα1^71,73–75^. In this work, we have shown that AMPKα2 is sufficient for metabolic stress-induced nuclear AMPK activity. Input-dependent AMPKα isoform specificity could provide another level of control over nuclear AMPK activity. These additional regulatory controls would ensure precise tuning of nuclear AMPK activity and allow the kinase to discriminate between downstream effectors.

In summary, we have generated a new single-fluorophore excitation-ratiometric AMPK activity reporter, which we have used to uncover mechanisms of compartmentalized AMPK activity. Using our new reporter, precise and sensitive investigation into spatial AMPK signaling is now possible, which should lead to a better understanding of metabolic regulation throughout the cell.

## Author Contributions

D.L.S. and J.Z. conceived the project; D.L.S., S.D.C., S.M., R.J.S., and J.Z., designed experiments, D.L.S, J.F.Z, M.C, S.D.C., and C.Y.H. performed experiments and analyzed data. A.L. and P.R. performed mathematical modeling. J.Z., R.J.S., and P.R. coordinated the study and provided guidance. D.L.S., S.M., and J.Z. wrote the paper. All authors discussed the results and approved the final version of the manuscript.

## Acknowledgements

We thank the following: Brian Tenner, Megan Mizinski, and Daniel Nguyen for assistance with molecular cloning; Qiang Ni for support in material acquisition; Eric Greenwald for assisting with MATLAB code development; Eric Griffis and Daphne Bindels with the UC San Diego Nikon Imaging Center; and Xin Zhou and Anne Lyons for helpful discussion and critique. This work was supported by the National Institutes of Health (NIH/NCI T32 CA009523 and NIH/NIGMS K12 GM068524-17 to D.L.S.; R35 CA197622, R01 DK073368, and R01 DE030497 to J.Z.; and R35 CA220538 and P01 CA120964 to R.J.S.); the University of California President’s Postdoctoral Fellowship (to D.L.S.); and the Air Force Office of Scientific Research (FA9500-18-1-0051 to J.Z. and P.R.).

## Experimental Procedures

### Materials

2-deoxyglucose (2-DG, Sigma, D6134-250MG) was dissolved in 1x DPBS. SBI-0206965 (Cayman Chemical, 18477), MK-8722 (Aobious, AOB33226), and CCCP (Fisher Scientific, AC228131000) were dissolved in DMSO (Sigma Aldrich). Hoescht 33342 (Cell Signaling, 4082S) was dissolved in deionized water. For western blotting and immunostaining, antibodies from Cell Signaling Technologies (Denvers, MA USA) were used at 1:1000 dilution unless otherwise noted: P-ACC S79 (3661), P-AMPK T172 (2535), ACC (3662), AMPKα (2532), GAPDH (5174, 1:10,000), Laminin A/C (4777), β-tubulin (2146, 1:10,000). From Sigma-Aldrich, antiActin (A5441) was diluted 1:10000 and anti-Tubulin (T5168) was diluted 1:5000. Lipofectamine 2000 (11668019) was purchased from ThermoFisher, and FuGENE HD (E2311) was purchased from Promega. MitoTracker Red (M22425) and LysoTracker Red(L7528) were purchased from ThermoFisher and diluted in DMSO.

### Plasmids

All primers used for molecular cloning can be found in Supplemental Table 4. To generate ExRai AMPKAR, DNA fragments encoding cpGFP and FHA1 binding domain and linker pair candidates obtained for our protein kinase A sensor ExRai AKAR^21^ were digested with SacI and EcoRI restriction enzymes (ThermoFisher, FD1134 and FD0275, respectively) and ligated into a SacI/EcoRI-digested pRSET-B backbone containing an AMPK substrate sequence. These constructs were then subcloned into the pcDNA3 vector via digestion with BamHI (ThermoFisher FD1464) and EcoRI restriction enzymes. ExRai AMPKAR T/A was generated via Gibson Assembly using NEBuilder HiFi DNA Assembly Kit (New England Biolabs E2621) using primers 1-2. Mitochondrial (MAIQLRSLFPLALPGMLALLGWWWFFSRKKADP), and nuclear (PKKKRKVEDA) ExRai AMPKAR were made by PCR-amplifying ExRai AMPKAR using primers 3-6 followed by insertion into BamHI/EcoRI-digested vector backbones containing the indicated localization sequences. Lysosomal ExRai AMPKAR was made by inserting PCR-amplified LAMP1^49^ generated using primers 7-8 into HindIII (ThermoFisher FD0504)/BamHI-digested ExRai AMPKAR backbone. mScarlet-AMPKα2 and 3x NLS-mScarlet-AMPKα2 (NLS targeting: PKKKRKVDPKKKRKVDPKKKRKV) in pEGFP-N1 expression vector were generated via Gibson Assembly using primers 9-14. AMPKα2 mutants were generated using primers 15-18. Successful clone generation was confirmed by Sanger sequencing (Genewiz). ABKAR^18^ and mCherry-LKB1^18^ were previously reported. EGFP-AMPKα2^57^ was a gift from Jay Brenman (Addgene plasmid 30310). H2B-mCherry was a gift from Michael Davidson (Addgene plasmid 55055).

### Cell Culture and Transfection

HEK293T cells were acquired from ATCC and cultured in Dulbecco’s modified Eagle medium (DMEM; Gibco 11885-084) containing 1 g/L glucose, 10% fetal bovine serum (FBS, Gibco 26140-079), and 1% (v/v) penicillin-streptomycin (Pen/Strep, Gibco 15140-122). WT MEFs, AMPK KO MEFs, and LKB1 KO MEFs were described previously^28,39^ and cultured in DMEM (Gibco 11995-065) containing 4.5 g/L glucose, pyruvate, and L-glutamine, supplemented with 10% FBS and 1% Pen-Strep. Cos7 cells were obtained from ATCC and cultured under the same conditions as MEFs. All cells were grown in humidified incubators kept at 37C and 5% CO_2_ (HeraCell) and checked for mycoplasma using Hoechst staining. For transfection, cells were plated on 35mm glass-bottomed dishes (CellVis D35-14-1.5-N). Cells were transfected 2-24 hours after plating. HEK293T cells and Cos7 cells were transfected with DNA using Lipofectamine 2000, and MEFs were transfected with DNA using FuGENE HD. Cells were imaged 24-48 hours after transfection.

### Fluorescence Imaging and Image Analysis

For all live cell imaging experiments, cells were washed and incubated in Hanks balanced salt solution (HBSS, Gibco 14065-056; buffered with 20 mM HEPES, pH 7.4 and supplemented with 2 g/L glucose) for at least 30 min at 37°C prior to imaging. All imaging was performed in the dark at 37°C. DMSO (1-3 uL/mL), 2-DG (40 mM), MK-8722 (500 nM), SBI-0206965 (30 μM), and CCCP (5 μM) were added at the indicated times.

Time-lapse epifluorescence images were acquired on either a Zeiss AxioObserver Z1 microscope (Carl Zeiss) equipped with a plan-apochromat 20x/0.8 N/A and 40x/1.4 N/A objectives (Carl Zeiss) and CMOS Orca Flash 4.0 camera (Hamamatsu) enclosed in a custom incubator, or a Zeiss AxioObserver Z7 microscope equipped with a 40x/1.4 N/A objective, Prime 95B sCMOS camera (Photometrics), and stage-top incubator (Carl Zeiss). Imaging experiments were done using a modified version of the open-source MATLAB (Mathworks) and μmanager (Micro-Manager)-based MATScope imaging suite (GitHub). Dual GFP excitation-ratio imaging was accomplished using ET405/40x and ET480/30x excitation filters with a T505dcxr dichroic, and a ET535/50m emission filter. EGFP was imaged using an ET480/30x excitation filter with a T505dcxr dichroic, and ET535/50m emission filter. mCherry and mScarlet were imaged using an HQ568/55x excitation filter with a Q600LPxr dichroic and HQ653/95m emission filter. Dual cyan/yellow emission ratio imaging was performed using an ET420/20x excitation filter, a T4551pxt dichroic, and two emission filters (ET470/40m for CFP and ET535/25m for YFP). CFP imaging was done using ET420/20x excitation filter with a T4551pxt dichroic and AT470/40m emission filter. YFP imaging was done using ET495/10x excitation filter with a T5151p dichroic and ET535/25m emission filter. All filter sets were controlled by an external filter-exchanger (Prior Scientific or Ludl Electronic Products, Ltd). Exposure times ranged between 50 to 100 ms, and images were acquired every 15-60 s.

Image analysis for time-lapse imaging was done using custom MATLAB code. Regions of interest (ROI) were randomly selected in cells throughout the field of view. For localized biosensors, ROIs were selected around the mitochondria, a region of the cell containing lysosomes, or the entire nucleus. Raw fluorescence intensities were corrected for background fluorescence using a cell-free area. cpGFP excitation ratios (Ex488/400) and yellow/cyan FRET emission ratios were calculated for each time point. Ratios were normalized to values before drug stimulation. Maximum ratio changes (ΔR/R_0_) were calculated as (R_max_-R_0_)/R_0_, where R is the excitation or emission ratio. SNR for each cell was calculated by dividing the maximum ratio change by the standard deviation of the baseline before drug addition. Z-factors were calculated as 1 – (3σ_max response average_ +3σ_baseline average_)/| σ_max response average_ - σ_baseline average_|. Graphs were plotted using GraphPad Prism 8 or later (GraphPad Software).

Fluorescence recovery after photobleaching experiments were performed on a Nikon Ti2 C2 confocal microscope equipped with a 40x 1.3 NA oil objective (Nikon), 405 nm, 488 nm, and 561 nm laser lines, C2-DUVB GaAsP detector, and Okolabs stagetop incubator. The microscope was controlled using NIS Elements. Nuclei were identified by H2B-mCherry expression. The entire nucleus was selected for photobleaching. The nucleus was photobleached for 1 second with 100% 488 nm laser power and observed for at least 25 min post-bleaching. Unnormalized intensity data was collected using NIS Elements and normalized using Excel (Microsoft). FRAP curves were fit to a single exponential recovery equation using the curve fitting tool in FIJI^76^, *y*=*a*(*1-e*^-bx^)+*c*.

### Western Blotting

MEFs were plated in either 6-cm or 10-cm dishes in growth medium. Cells washed with HBSS and treated with small molecules. After drug treatment, cells were scraped on ice and pelleted in PBS. PBS was aspirated and the cells where lysed in lysis buffer (3x radioimmunoprecipation assay buffer, protease inhibitor cocktail (Roche 1169749801), 1 mM phenylmethylsulfonyl fluoride, 1 mM sodium pervanadate, 1 mM sodium fluoride, and 25 mM calyculin A). Protein samples were separated using 4-15% SDS-PAGE (BioRad 456-1085) and transferred to nitrocellulose membranes. Membranes were blocked using 1% bovine serum albumin (SigmaAldrich 03116956001 in 1x TBS, 0.05% Tween-20) and incubated with primary antibodies overnight at 4°C. After incubation with the appropriate horseradish peroxidase-conjugated secondary antibody, bands were visualized by chemiluminescence.

### Nuclear Fractionation

MEFs were seeded at a density of 6×10^6^ cells per 15-cm dish in growth medium 18 hours prior to treatment. Small molecules were applied to cells and incubated for 60 min at 37°C: DMSO (3 μL/mL), 2-DG (40 mM), MK-8722 (500 nM). Cells were scraped on ice and pelleted in PBS. PBS was aspirated and cells were taken forward for nuclear/cytoplasmic fractionation using NE-PER kit (ThermoFisher Scientific, 78835) following manufacturer’s protocol, or for whole-cell lysate control samples. Fractionation was performed using WT MEFs and AMPK KO MEFs. Parallel whole-cell lysates were generated using CST lysis buffer (20mM Tris, pH 7.5, 150 mM NaCl, 1 mM EDTA, 1 mM EGTA, 1% Triton X-100, 2.5 mM pyrophosphate, 50 mM NaF, 5 mM β-glycero-phosphate, 50 nM calyculin A, 1 mM Na_3_VO_4_, and protease inhibitors). Lysates were incubated at 4°C for 15 min and cleared at 16,000 x g for 10 min at 4°C. Total protein was normalized using the BCA Protein Assay kit (Pierce 23225).

### Development of Mathematical Model of Mitochondrial AMPK Activity

The mathematical model was constructed using COPASI (Build 226). In this model, we consider only the effects of AMPK activation in the cytosol proximal to the mitochondrial membrane. We first consider the adenine nucleotide equilibrium, which is mediated by adenylate kinase^32^. Likewise, we model AMPK activation by adenine nucleotides as in Connolly^32^. To model the activation of AMPK biosensor, as well as AMPK activity, we derive equations from Durandau^31^. These equations describe the activity of a kinase given experimental measurements of a biosensor modeled as Michaelis-Menten activation. We use this equation to fit AMPK activity to biosensor measurements through parameters, V, R, and ***τ*** (Supplemental Tables 1-3).

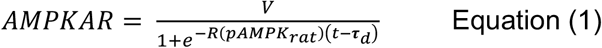

Finally, we model the metabolic fluxes of glycolysis, ATP hydrolysis, and oxidative phosphorylation as simple mass action fluxes, scaled to metabolic rates in MEFs. ATP hydrolysis is a free parameter in our model that was altered and led to a range of cellular behavior.

To model pharmacological inhibition of metabolic pathways, we altered the rates of glycolysis and oxidative phosphorylation at the time of 2-DG application. We then simulate the model and show changes in ATP concentration and AMPKAR activation.

### Statistics and Reproducibility

Figure preparation and statistical analysis were performed using GraphPad Prism 8 or later versions. For comparison of two parametric data sets, the student’s t-test was used. Nonparametric tests were done using Mann-Whitney U. For comparing three or more sets of data, ordinary one-way ANOVA followed by multiple comparisons test was done. Statistical significance was defined as P < 0.05 with a 95% confidence interval. The number of cells analyzed (n cell) and number of independent experiments are reported in all figure legends. All time courses shown are the mean of all cells ± standard deviation (SD) unless otherwise noted. All dot-plots shown depict the mean ± standard error of the mean (SEM).

## Data Availability

All data and materials supporting these findings are available upon reasonable request.

## Code Availability

Custom MATLAB code used for image analysis is available upon reasonable request.

## Notes

### Competing Interest Statement

The authors have declared no competing interest.

