## Supplemental Figures and Legends for "Spatial regulation of AMPK signaling revealed by a sensitive kinase activity reporter"

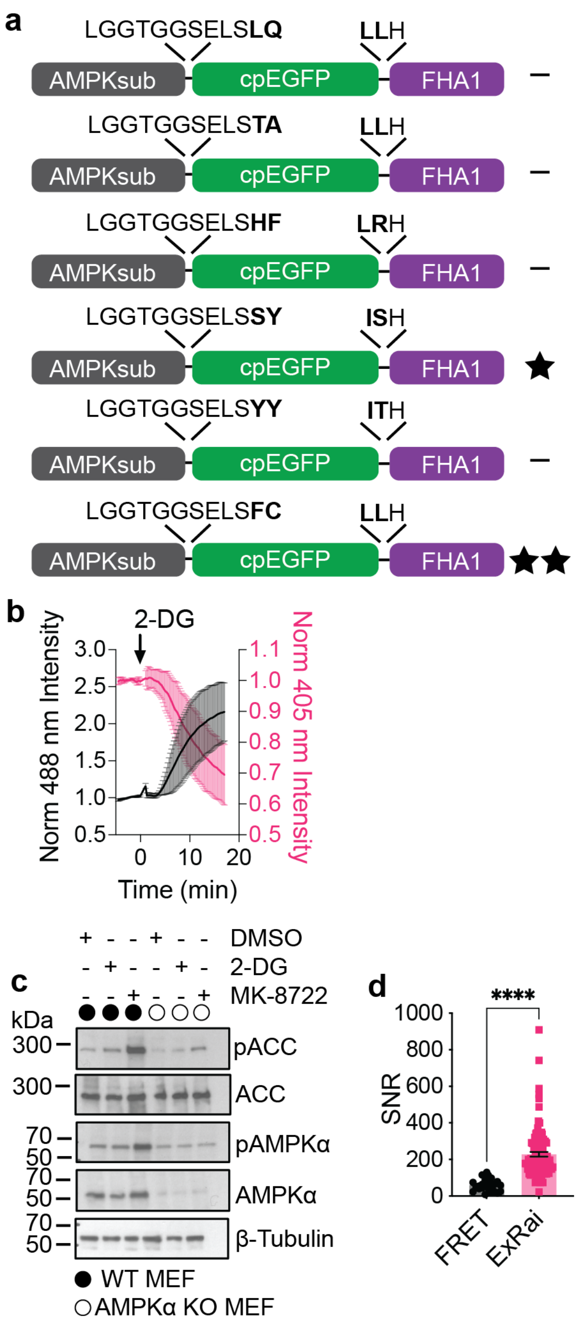


Schmitt et al Supplemental Figure 1

**Supplemental Fig 1. Development and characterization of ExRai AMPKAR.**

**a,** Domain structures of ExRai AMPKAR linker variants tested. Variants are scored based on performance: -, <40%; ★, 40-120%; ★★, >120% change in excitation ratio.

**b**, Average normalized 480 nm-excited fluorescence intensity (black), 405 nm-excited fluorescence intensity (pink) of Cos7 cells from Fig 1b expressing ExRai AMPKAR in Cos7 cells treated with 2-DG (40 mM).

**c**, Western blot of AMPK activity in WT and AMPK KO MEFs treated with DMSO, 2-DG (40 mM), or MK-8722 (500 nM) for 30 min. Western blots are representative of at least 3 replicates.

**d**, Signal-to-noise ratio (SNR) of ABKAR (black) and ExRai AMPKAR (pink, ****$P<0.0001$, unpaired t-test).

Time courses show the mean ± SD, dot plots are shown as mean ± SEM.

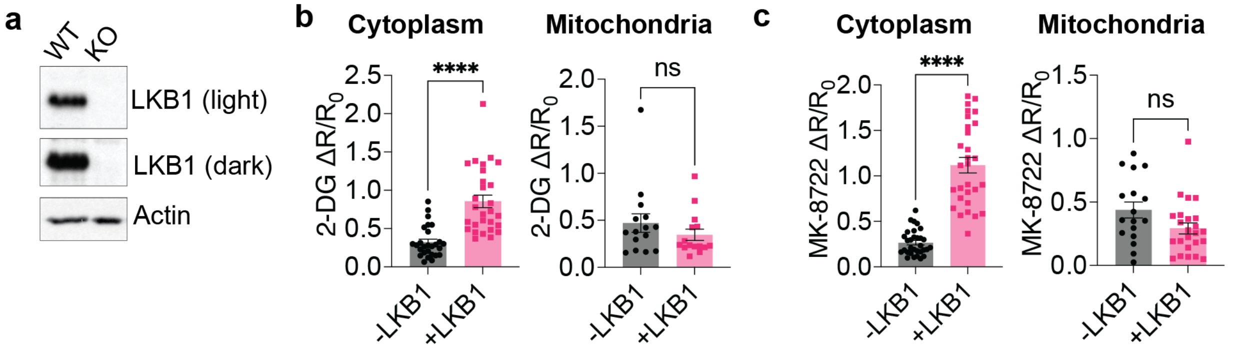


Schmitt et al Supplemental Figure 2

**Supplemental Figure 2. Lysosomal AMPK activity measured by ExRai AMPKAR.**

**a**, Western blot of LKB1 expression in WT and LKB1-KO MEFs, representative of at least two experiments.

**b**, (Left) Maximum ExRai AMPKAR ratio change in HeLa cells without (black, n = 27 cells from 3 experiments) or with (pink, n = 28 cells from 3 experiments) mCherry-LKB1 expression in response to 2-DG (40 mM; ****P<0.0001, unpaired t-test). (Right) Maximum ratio change of mito-ExRai AMPKAR in HeLa cells without (black, n = 15 cells from 3 experiments) or with (pink, n = 15 cells from 3 experiments) mCherry-LKB1 expression in response to 2-DG (40 mM; P = 0.25, unpaired t-test).

**c**, (Left) Maximum ratio change of cytoplasmic ExRai AMPKAR in HeLa cells without (black, n = 31 cells from 3 experiments) or with (pink, n = 29 cells from 3 experiments) mCherry-LKB1 expression in response to MK-8722 (500 nM; ****P<0.0001, unpaired t-test). (Right) Maximum ratio change of mito-ExRai AMPKAR in HeLa cells without (black, n = 17 cells from 3 experiments) or with (pink, n = 24 cells from 3 experiments) mCherry-LKB1 expression in response to MK-8722 (500 nM; P = 0.056, unpaired t-test). Dot plots show the mean ± SEM.

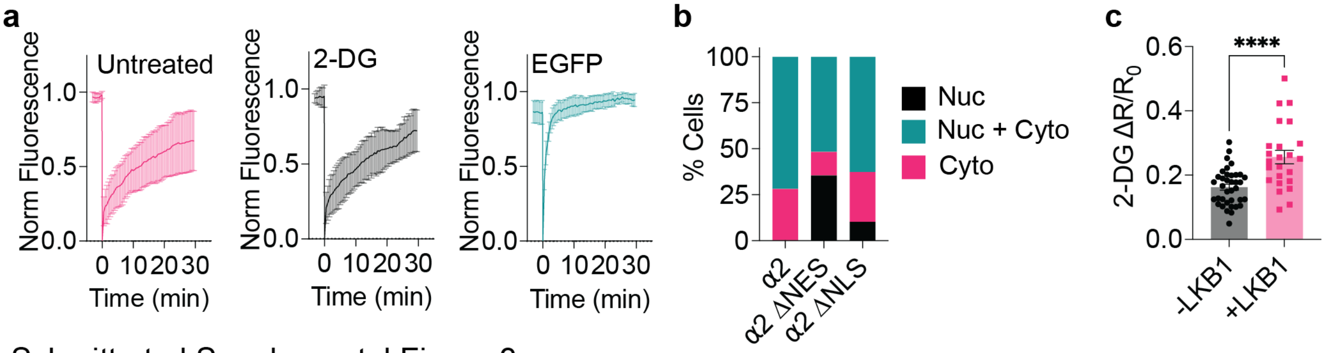


Schmitt et al Supplemental Figure 3

**Supplemental Figure 3. Nuclear AMPK activity is due to shuttling of AMPKα2 from the cytoplasm to the nucleus.**

**a**, Average FRAP recovery curves of nuclear EGFP-AMPKα2 in AMPKα KO MEFs treated with either DMSO or 2-DG (40 mM) immediately before FRAP experiment began, as well as EGFP control.

**b**, Quantification of mScarlet-AMPKα2, mScarlet-AMPKα2 ΔNES (L546A and L550A), mScarlet-AMPKα2 ΔNLS (K224A) localization to either only the nucleus (nuc), cytoplasm (cyto), or nucleus and cytoplasm (nuc + cyto). Representative of two independent experiments.

**c**, Average response of nuclear-ExRai AMPKAR in HeLa cells without (black; n =37 cells from 4 experiments) and with (pink; n = 24 cells from 4 experiments) mCherry-LKB1 expression and treated with 2-DG (40 mM) (****P < 0.0001). Response curves show the mean ± SD, dot plots show the mean ± SEM.

### Supplemental Table 1. AMPK Activity Equation Development

$$\text{AMPKAR} = \frac{V}{1 + e^{-R(pAMPKrat)(t-\tau_d)}}$$

| Parameter | Mean | St. Dev. |
|---|---|---|
| V | $0.145 \times 10^{-2}$ | $0.180 \times 10^{-2}$ |
| R | $0.795 \times 10^{-2}$ | $0.651 \times 10^{-2}$ |
| $\tau_d$ | 920.6 | 265.6 |

### Supplemental Table 2. Table of Reactions, Fluxes, and Parameters

| Reaction | Flux Expression | Parameters | References |
|---|---|---|---|
| $ATP + AMP = ADP + ADP$ | $J_{AEQ} = k_{f,AEQ}[ATP][AMP] - k_{r,AEQ}[ADP][ADP]$ | $k_f = 7.3 \times 10^{-5}[1/(\mu Ms)]$<br>$k_r = 4.5 * 10^{-5}[1/(\mu Ms)]$ | 2 |
| $ATP \rightarrow ADP$ | $J_{HY\,D} = k_{HY\,D}[ATP]$ | $k_{HY\,D} = 1 \times 10^{-5}[1/(\mu Ms)]$ | 1, 2, 3 |
| $AMP + AMPK \rightarrow ATP + pAMPK$ | $J_{AMPK} = k_{f,AMPK}[AMP][AMPK]$ | $k_f = 1.3[1/(\mu Ms)]$ | 2 |
| $ATP + pAMPK \rightarrow ATP + AMPK$ | $J_{AMPK} = k_{f,AMPK_r}[ATP][pAMPK]$ | $k_f = 0.2[1/(\mu Ms)]$ | 2 |
| $ADP \rightarrow ATP$ | $J_{GLYC} = k_{GLYC}[ADP]$ | $k_{GLYC} = 6.602 \times 10^{-5}[1/(\mu Ms)]$ | |
| $pyruvate + ADP \rightarrow ATP$ | $J_{OP} = k_{OP}[ADP]$ | $k_{OP} = 0.001[1/(\mu Ms)]$ | |


### Supplemental Table 3. Table of Equations, Initial Conditions, and Parameters

| Species | Differential Equations | Initial Conditions $[\mu M]$ |
|---|---|---|
| AMP | $\frac{d[AMP]}{dt} = -J_{AEQ} - J_{AMPK}$ | 27 |
| ADP | $\frac{d[ADP]}{dt} = +J_{AEQ} - J_{GLYC} - J_{OP} + J_{HY\,D}$ | 270 |
| ATP | $\frac{d[ATP]}{dt} = J_{AEQ} + J_{AMPK}{}_{J_{GLYC}} + J_{OP} - J_{HY\,D}$ | 2700 |
| AMPK | $\frac{d[AMPK]}{dt} = -J_{AMPK}$ | 0.144 |
| pAMPK | $\frac{d[pAMPK]}{dt} = +J_{AMPK}$ | 0.081 |
| pAMPKAR | $AMPKAR = \frac{V}{1 + e^{-R(pAMPKrat)(t-\tau_d)}}$ | 1 [1] |

**Supplemental Table 4. Primers used for molecular cloning**

| Primer Number | Purpose | Sequence (5' to 3') |
|---|---|---|
| 1 | ExRai AMPKAR T/A, forward | GACGATAAGGATCCCATGAGGAGAGTGGCTGCTCTGGTGGATCTGGGC |
| 2 | ExRai AMPKAR T/A, reverse | CCGCCAGTGTGATGGATATCTGCAGAATTCCTAGCGATCAACTTTGTTCTGCTCGAGGCA |
| 3 | ExRai AMPKAR, forward | AAGGATCCCATGAGGAGAGTGGCTACCCTG |
| 4 | ExRai AMPKAR, reverse with stop codon | CCGCCAGTGTGATGGATATCTGCAGAATTCCTAGCGATCAACTTTGTTCTGCTCGAGGCA |
| 5 | ExRai AMPKAR, to remove stop codon forward | CAGAACAAAGTTGATCGCTATGAATTCCCCAAAAAGAAG |
| 6 | ExRai AMPKAR, to remove stop codon reverse | CTTCTTTTTGGGGAATTCATAGCGATCAACTTTGTTCTG |
| 7 | LAMP1, forward | GCGCAAGCTTGCGGCCGCCACCATGGCGGCGCCCGGCAGCGCCCGG |
| 8 | LAMP1, reverse | GCGCGGATCCGCACCACCGCCACCACCGATAGTCTGGTAGCCTGCG |
| 9 | mScarlet-AMPKa2, forward | TCAGATCCGCTAGCGCTACCGGTCGCCACCATGGTGAGCAAGGGCGAGGCAGTGATCAAG |
| 10 | mScarlet-AMPKa2, reverse | GCTTGAGCTCGAGATCTGAGTCCGGCCGGACTTGTACAGCTCGTCCATGCCGCCGGTGGA |
| 11 | 3x NLS-mScarlet-AMPKa2, forward | AGAAAGGTAGGATCCAGTATGGTGAGCAAGGGCGAGGCAGTGATCAAG |
| 12 | 3x NLS-mScarlet-AMPKa2, reverse | CTTGATCACTGCCTCGCCCTTGCTCACCATACTGGATCCTACCTTTCT |
| 13 | 3x NES-mScarlet-AMPKa2, forward | GAACGACTGACCCTGGATATGGTGAGCAAGGGCGAGGCAGTGATCAAG |
| 14 | 3x NES-mScarlet-AMPKa2, reverse | CTGCCTCGCCCTTGCTCACCATATCCAGGGTCAGTCGTTCCAGTGGGGGCAGATCCAGGG |
| 15 | AMPKα2 ΔNLS | AAGCCCAAATCTTTAGCTGTGAAAGCCGCCAAGTGGCACCTTGGGATCCG |
| 16 | AMPKα2 ΔNES | GAAATGTGCGCCAGTGCAATCACTGCTGCAGCCCGTTGAGGATCC |
